# *In Silico* Structure-Based Interactomic Analysis of the Scaffolding Protein DCAF7

**DOI:** 10.64898/2026.05.13.724911

**Authors:** Alexandre Mezghrani, Victor Reys, Gilles Labesse

## Abstract

WD40 domains share a widespread β-propeller fold, and often act as versatile scaffold proteins. Despite their central role in organizing dynamic cellular complexes, the molecular and structural mechanisms of many WD40 proteins remain poorly understood. Among them, DCAF7, an ubiquitously expressed and essential gene in human, also encodes a highly conserved WD40 protein in eukaryotic organisms. It is known to interact with multiple and functionnally diverse partners to coordinates cellular activity of several protein kinases as well as transcriptional regulators, thereby modulating key cellular processes such as cell growth, differentiation, and transcriptional regulation. However, the precise mode of action of DCAF7 is unknown and its important divergence in sequence from better characterize WD40 prevent information transfer by similarity.

Structural interactomic can reveal how protein–protein interactions (PPIs) occur within an organism and are essential for understanding biological functions and developing new therapeutic strategies. Using SLiMAn2, AlphaFold2/3 and PSSMsearch, we identified a conserved α-helical short linear motif (SLiM) in several well known DCAF7 partners that binds to the top surface of its β-propeller. This motif was subsequently used to generate a regular expression, to identify potential new direct binders across the DCAF7 meta-interactome and the human proteome. Domain-domain interactions were also predicted for some other partners. Finally, modeling of oligomeric complexes with such new hits reveals the structural basis of DCAF7 scaffolding, with links to neurodevelopmental disorders such as autism.

## Introduction

Among the most abundant domain families in the human proteome, WD-repeat or WD40 domains show remarkable diversity in both molecular features and biological functions (Schapira et al. 2017; Ackloo et al. 2024). This diversity limits the generalization of conserved functions within the family due to high sequence divergence. The WD40 domain adopts a β-propeller fold composed of seven blades arranged radially, forming a donut-like shape. This architecture provides multiple potential interaction surfaces, increasing the versatility of WD40 proteins in engaging diverse partners (Jain et Pandey 2018; Schapira et al. 2017). Several WD40 proteins bind partners via SLiMs at the top of the β-propeller, but can also have PPIs interfaces at the side (inter-blade) or bottom surfaces (Gurung et al. 2023; Cao et al. 2014; Fukumoto et al. 2008). A significant fraction (30 to 40%) of human WD40 proteins are poorly characterized experimentally at the structural level, although accurate AlphaFold models have been generated.

An example is the highly conserved WD40 protein DCAF7 (DDB1- and CUL4-associated factor 7), also known as HAN11 or WDR68, which interacts with a range of diverse proteins (Morriss et al. 2013; Yang et al. 2006; Glenewinkel et al. 2016), and the corresponding binding interfaces are yet to be precisely delineated.

DCAF7 is mainly known to interact with the kinases DYRK1A and DYRK1B, which are involved in the regulation of cell cycle progression, transcription factor activity, and developmental signaling (Skurat et Dietrich 2004; Aranda et al. 2011). These interactions are essential for proper cellular function, as they allow DYRK1A and DYRK1B to be stabilized and appropriately localized (Ritterhoff et al. 2010; Yousefelahiyeh et al. 2018). In addition, the coordinate binding of M3K1, HIPK2 and DYRK1A kinases to DCAF7 forms a scaffolded kinase complex that coordinates phosphorylation of shared substrates, integrating developmental and stress-response signaling pathways (Ritterhoff et al. 2010). DCAF7 modulates HIPK2 ability to phosphorylate downstream substrates such as p53 and components of the TGF-β pathway (Ritterhoff et al. 2010). Through these interactions, DCAF7 contributes to the fine-tuning of cellular responses, linking kinase-mediated phosphorylation with transcriptional and post-transcriptional regulation, thereby ensuring coordinated execution of complex biological programs.

A growing body of evidence indicates that DCAF7 is also present in protein complexes involved in neurodevelopmental processes. DCAF7 is an integral component of a non-canonical Polycomb Repressive Complex 1 (PRC1-AUTS2) that includes AUTS2, PCGF5, RING1A/B, RYBP/YAF2 and Casein kinase 2. Within this complex, DCAF7 is required for PRC1-AUTS2-mediated transcriptional activation of genes involved in neuronal differentiation (Gao et al. 2014; Wang et al. 2018; Geng et Gao 2020). Its deletion in mouse embryonic stem cells leads to defects in neuronal lineage commitment and down-regulation of key neurogenic transcripts (Gao et al. 2014; Wang et al. 2018). DCAF7 knockout exhibit intrauterine growth retardation in mouse embryos and impaired neural tube development and neurogenesis (Wu et al. 2025; Nissen et al. 2006)). DCAF7 has been also shown to be link with the cytoplasmic protein Huntingtin-associated protein 1 (Hap1), particularly in hypothalamic neurons where these proteins co-localize in stigmoid bodies (Xiang et al. 2017). Hap1 and DYRK1A compete for DCAF7 binding, and the reduced Hap1-DCAF7 interaction in Down syndrome models correlates with delayed growth and altered neural development (Xiang et al. 2017).

DCAF7 has also been characterized as a component of CRL4 E3 ubiquitin ligase complexes through its association with DDB1 and CUL4, regulating the stability of several proteins involved in DNA repair and proteostasis (Higa et al. 2006; Xu et al. 2023; Yu et al. 2019; Peng et al. 2016). Among these are DNA ligase I and the ERCC1-XPF endonuclease, where DCAF7-dependent ubiquitination regulates cellular responses to DNA damage (Peng et al. 2016; Xu et al. 2023).

Although DCAF7 has essential roles in multiple cellular pathways, the molecular and structural basis of its pleiotropic functions are still poorly understood. This lack of mechanistic insight limits our understanding of how DCAF7 mediates its scafolding functions meanwhile several partners (such as DYRK1A and AUTS2) points to a putative role in human diseases, particularly neurodevelopmental disorders such as autism (Bicks et al. 2023; Earl et al. 2017; Oksenberg et al. 2013).

Partners of WD40 proteins may interact through domain-domain as well as via short linear motifs (SLiMs) interfaces (Davey et al., 2023). PPIs mediated by SLIMs are typically more transient and dynamic than domain-domain contacts and play key roles in signaling, regulation, and the assembly of multiprotein complexes (Oldfield & Dunker, 2014; Uversky, 2016). SLiMs are normally difficult to distinguish from non-interacting segments using AlphaFold confidence metrics (Bret et al. 2024). Nevertheless, successful strategies have been developed to increase the accuracy of AF for *de novo* identification of SLiMs (Bret et al. 2024; Lee et al. 2024; Veinstein et al. 2025). For the identification of SLiMs in a defined meta-interactome, the SLiMAn2 webserver is a well suited tool (Reys et al. 2024). It integrates several interactomic and structural metrics to prioritize functional SLiM occurrences, including the number of experimentally curated interactors (BioGRID, IntAct), predictions of SLiM disorder, the presence of post-translational modifications (PTMs), and links to literature co-occurrence (Reys et al. 2024 ; Mezghrani et al. 2024). Here, we used AF in combination with SLiMAn2, in an attempt to characterize conserved SLiMs present in several proteins within the DCAF7 meta-interactome. In aaddition, PSSMsearch allowed us to extend further our search and to precise the definition of a DCAF7-binding signature.

The prediction by AF modeling of similar binding modalities among several DCAF7 partners allowed us to design putative SLiM signatures for DCAF7 binding and then to build a structural interactome. Several regular expressions were derived with PSSMsearch and used to interrogate the DCAF7 interactome and then the whole human proteome, revealing numerous additional candidate binders. We further showed by combining structural confidence from AF with network-based interconnectivity scores, that these predicted new DCAF7 partners have similar connectivity levels than known direct binders of the DCAF7 meta-interactome. Finally, the significant enrichment in autism-associated genes across both known and newly predicted interactors further supports the biological relevance of our predictive models. Finally, modelling of macromolecular assemblies gives a structural basis for the scaffolding function of DCAF7. These assemblies link signaling proteins implicated in neurodevelopmental disorders. In particular, the models suggest that DCAF7 potentially connect the kinase DYRK1A with synaptic proteins such as SHANK3 and components of the AMPA glutamate receptors, offering a possible structural and molecular explanation for biological processes associated with autism-related signaling networks. These findings provide a comprehensive view of DCAF7 as a central scaffold that integrates kinase signaling, ubiquitin-mediated regulation, and epigenetic control.

## Results

### In silico identification of SLiM motif recognizes by DCAF7

As modelled by AF, DCAF7 adopts a seven-bladed β-propeller structure (pTM=0.92) composed of seven WD40 repeats (Figure 1A). This organization creates two distinct faces, the top and the bottom (Figure 1B). In DCAF7, the top face is more evolutionary conserved PPI than the bottom one that exhibits greater sequence variability (Figure 1B). Another highly conserved region in DCAF7 extends from the top toward the interblade 3–4 side of the donut-shaped structure (figure 1B).

**Figure 1:**
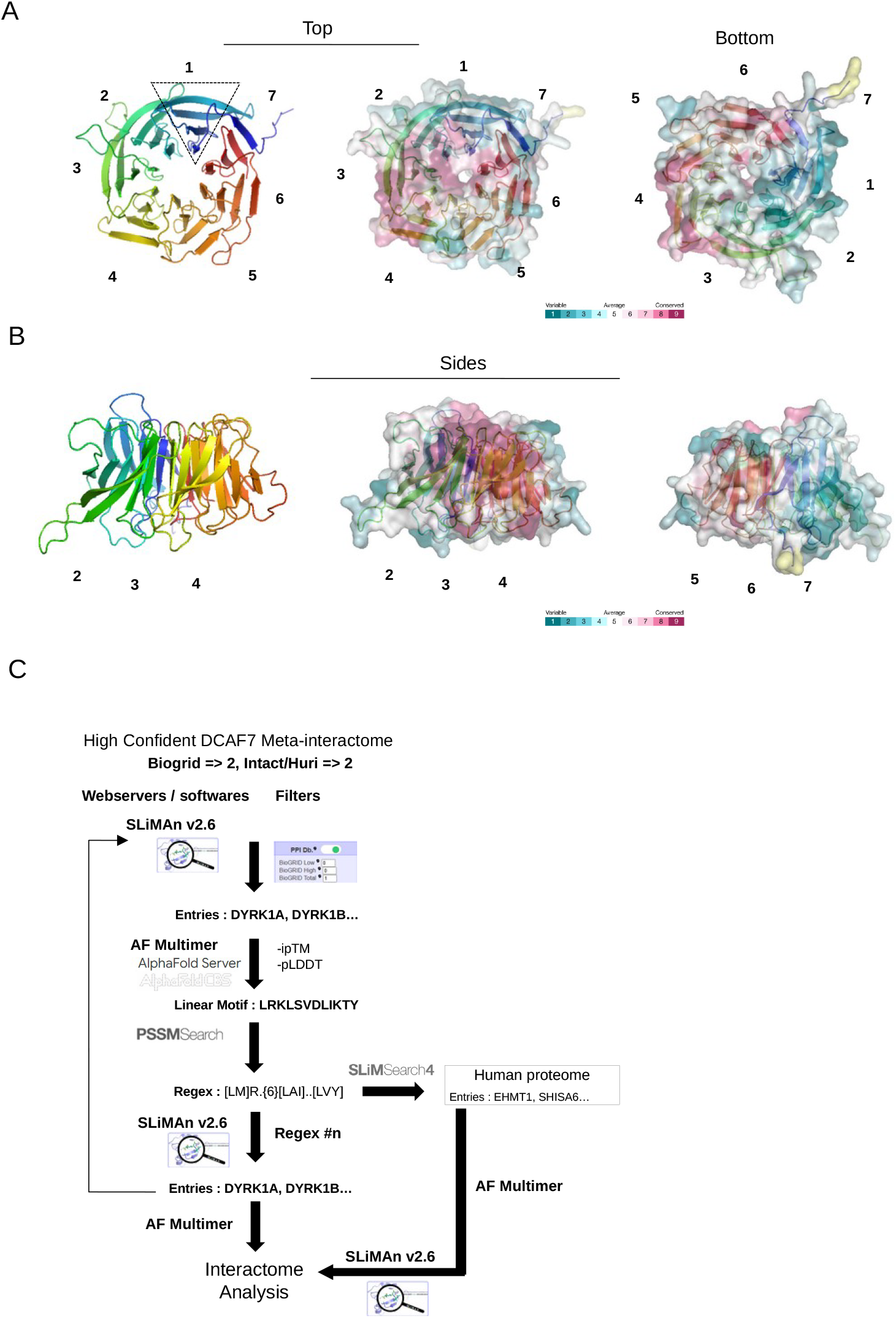
Strategy for *in silico* identification of SLiMs recognizes by the WD40 DCAF7. **A)** AF model of DCAF7 showing a β-propeller fold and highlighting the distinct regions. The WD40 repeats forming the seven blades (labeled 1–7) are colored for clarity. **B)** Top, bottom, and side views of DCAF7 surface conservation generated with ConSurf. Conservation is color-graded as indicated. **C)** Workflow used to identify conserved SLiMs recognized by DCAF7. Known DCAF7 partners were retrieved from BioGRID and IntAct and filtered for confidence using SLiMAn. Heterodimers were modeled with AF, and high quality models were retained for linear motif extraction. A regex is constructed using PSSMSearch and reintroduced into SLiMAn. Potential new interactors in the human proteome are identified using SLiMSearch4, and hits supported by HC AF models are selected for interactome analysis.

Although no SLiM motif is annotated for DCAF7 in the ELM database, we nevertheless used SLiMAn2, dedicated to the management and prediction of SLiMs, to extract and navigate the DCAF7 meta-interactome from BioGRID and IntAct datasets. SLiMAn2 generates a ranked list of 372 interactors, ordered from the most frequently to the least frequently observed in association with DCAF7. The list was, first, reduced to focus on the most confident partners of DCAF7, by retaining only the 55 proteins with at least two experimental evidences in both BioGRID and IntAct, and called this subset the high-confidence (HC) DCAF7 meta-interactome. We then structurally characterize their PPI interfaces with DCAF7 using AlphaFold2 multimer and AlphaFold3 (AF) (Figure 1C)(Alderson et al. 2023; Abramson et al. 2024). The structural models were evaluated using TM-score (ipTM) and pLDDT confidence metric at the binding interfaces (pLDDTi), providing a robust assessment of binding interface reliability. For some of the best predicted heterodimer, a linear interface of the partner, displaying a pLDDTi > 80, was extracted and used to generate a regular expression (regex) for position-specific scoring matrix analysis using PSSMsearch (Krystkowiak et al. 2018). This signature was subsequently introduced back into SLiMAn2 to assess the structural and biological relevance of the predicted PPI motif in DCAF7 interactome. This stepwise approach is based on a hierarchical survey of the best characterized binders down to the more putative ones.

The most frequently detected partners of DCAF7 are the homologous kinases DYRK1A and DYRK1B. Their association with DCAF7 has been reported in both high-throughput and low-throughput experiments (Yousefelahiyeh et al. 2018; Glenewinkel et al. 2016). Moreover, a short binding motif has been deduced through co-IP experiments (Glenewinkel et al. 2016). Among the different AF models of the complex DCAF7-DYRK1A we generated, one linear motif in the N-terminal IDR of DYRK1A is invariably positioned in a similar manner within a conserved pocket at the top of the DCAF7 β-propeller (Figure 2A). This motif nicely matches the region identified experimentally. This binding motif in DYRK1A corresponds to a kinked α-helix (Pro92–Gly122), overlapping with the N-terminal autophosphorylation accessory region (NAPA). It is separated from the DH box (residues 137–147) and the kinase domain (140–470) by an unstructured segment (123–136) that includes a NLS (middle part, Figure 2A). The approximately 87-degree kink observed in the DYRK1A α-helix bound to DCAF7 is also present in the isolated monomer, indicating that is an intrinsic structural feature rather than one induced upon binding. However, some reorientations of the side chains of the residues Leu93, Arg94, Lys102, and Tyr104 are observed in DYRK1A bound to DCAF7 compared to monomeric DYRK1A alone (Figure 2B). Since the main DCAF7-binding domain in DYRK1A lies within an IDR, we also modeled 50- and 30-amino-acid fragments of this region, as previously proposed for SLiM detection (Bret et al. 2024). The resulting models showed high confidence scores, with ipTM and pLDDTi values of 0.94 and 90, respectively. Furthermore, the AF model using the 30-residue fragment shows perfect structural overlap (RMSD =0.34) with the binding interface of the full-length (FL) DYRK1A (Figure 2B).

**Figure 2:**
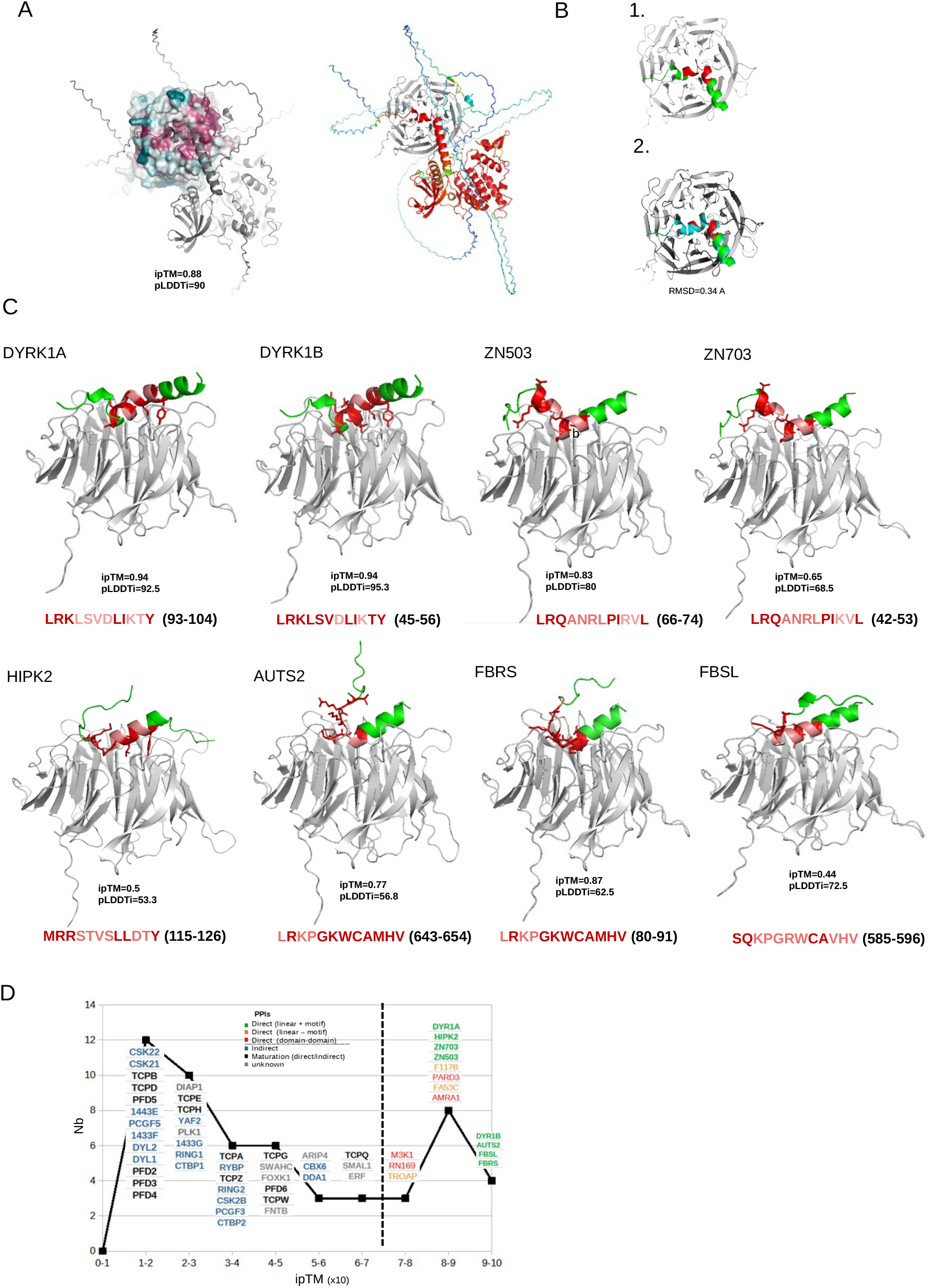
Structural modeling of conserved linear motif-based interfaces of DCAF7. **A)** AF model of DCAF7 in complex with DYRK1A: on the left, the surface conservation of DCAF7 is shown, colored by conservation grade (ConSurf, https://consurf.tau.ac.il/), with DYRK1A in cartoon view; on the right, DYRK1A is colored by pLDDT score and DCAF7 in cartoon view. **B)** 1) AF model of a 30-residue DYRK1A fragment bound to DCAF7. 2) The corresponding region of the full-length DYRK1A model was superimposed onto the fragment, and RMSD was calculated using PyMOL. **C)** AF models of 30-residue fragments of various DCAF7 partners. In each model, the fragment is shown in cartoon representation in complex with DCAF7, with the motif displayed below the model. Residues in contact with DCAF7 are highlighted in dark red. **D)** Repartition with AF ipTM scores of the HC DCAF7 interactors (55). Interactors with ipTM > 0,7 are considered direct (separated by a dashed line). The nature of the PPIs (direct vs. indirect) and the type of binding interfaces with DCAF7 are specified.

For DYRK1B, AF model consistently shows that DYRK1B engages DCAF7 through the same kinked α-helix (residues 45–56), and inserted in a similar manner into the conserved pocket at the top of the DCAF7 β-propeller (Figure 2C). The binding interface displays 100% sequence identity between DYRK1A and DYRK1B.

The third most frequent partner, the protein-kinase HIPK2 is functionally as well as physically associated with DYRK1A in several cell types (Ritterhoff et al. 2010; Glenewinkel et al. 2016). Despite a high ipTM score (0.82) of the FL HIPK2-DCAF7 heterodimer model, the binding domain of HIPK2 does not match the region experimentally defined (Glenewinkel et al. 2016). But, when considering the disordered regions of HIPK2 at both the N- and C-termini, the fragments 1–190 and 562–1198 display high ipTM scores (0.85 and 0.84, respectively). Interestingly, the experimentally defined motif within the N-terminal region (115–126) is positioned at the top of the β-propeller similarly to DYRK1A/B (Figure 2C). This motif adopts a straight helical conformation, unlike the kinked α-helix of DYRK1A/B (Figure 2C). AF modeling using 50- and 30-residue HIPK2 fragments centered on the DCAF7 motif also results in a SLiM-based interface. The pLDDTi for the corresponding HIPK2 fragments (55.9 for frag50 and 53.1 for frag30) is, however, relatively low (table 1). This might suggest either a lower affinity or a poor modeling of this particular interface.

Taken into account the first two motifs from DYRK1A and DYRK1B or adding on top of them the motif from HIPK2 led to the definition of two signatures using PSSMsearch. Including these signatures in SLiMAn analysis did not retrieve any additionnal partner from the HC DCAF7 meta-interactome, being likely too stringent (E. values of 1.9 × 10^−11^).

The next two most frequently interactors listed by SLiMAn, corresponding to the CK2 alpha subunits, did not show any reliable PPI interface with DCAF7 using AF, with very low ipTM score, 0.14 and 0.16 for CSK22 and CSK21 respectively. They may correspond to indirect interactions.

The sixth most frequently identified interactor of DCAF7, ZNF703, appears to fulfill the above structural criteria (supplementary table 1). The DCAF7–ZNF703 AF model predicted that a kinked α-helical peptide segment of ZNF703 (residues 42–53) binds within the conserved pocket at the top of the DCAF7 β-propeller, with a high pLDDTi score (88.6), as in the case of DYRK1A/B (Figure 2C). The peptide motif derived after addition of ZNF703 was subsequently used to define a new signature (regex #3: [LM]R.{6}[IL].[TV][YL]) with a less stringent E-value (3.7 × 10^−6^). This regex was also incorporated into SLiMAn, and retrieved one additional protein, ZNF503 in the DCAF7 HC meta-interactome. The DCAF7 binding motif of ZNF503 (residues 66–77) is highly similar to that of ZNF703, differing by only a single amino acid (R10K). A kinked helical structure is also predicted, further supporting the structural and functional conservation of this SLiM-based PPI (Figure 2C). Adding the ZNF503 motif sequence, resulted in an unchanged signature as derived by PSSMsearch. Although DCAF7–ZNF503/703 PPIs have been predominantly observed in high-throughput experiments and their molecular and structural basis remains largely unexplored, these findings provide some support for the validity of these new SLiMs and their functional relevance in relation to DCAF7 binding.

The next 4 proteins (TCPB, TCPA, M3K1, DIAP1) do not possess comparable SLiM-based interfaces with DCAF7 according to AF modeling and PSSMsearch. This suggested an alternative mode of binding or indirect interactions (see below).

Next, AUTS2, was identified with a very high ipTM (0.96) and pLDDTi (87) for its heterodimerization with DCAF7. Its AF model shows a helical motif for AUTS2, similar to that of HIPK2, binding within the same conserved pocket of DCAF7, suggesting a recurring SLiM-based binding mode in these interactors (Figure 2C). Addition of the AUTS2 motif sequence to PSSMsearch generates a new signature (regex #4:[LM]R.{6}[LAI].. [LVY]), with a much higher E-value (1.0 × 10^−4^) than the previous ones. This regex #4 retrieves 14 proteins in the HC meta-interactome. Among these, only one, FBRS, a AUTS2 paralog, exhibits the same binding mode as the previously described with high iptM (0.93) and pLDDTi (86) (table 1, Figure 2C). Next, FBSL, another AUTS2 paralog that is ranked immediately after AUTS2 among the most frequent DCAF7 binders and followed by FBRS displays the same PPI modality as AUTS2, with high ipTM (0.93) and pLDDTi (87). Incorporating the FBSL motif sequence allowed us to generate a new, more stringent regular expression (regex #5: [LMS][RQ][RKQ].{4} [LCP][ILA]..[LVY]), with a slightly better E-value (1.5 × 10^−5^). Beyond the 8 motifs used to construct this regular expression, no additional motif was identified with this regex among the HC DCAF7 interactors.

### Other direct binders within the HC meta-interacome

Next, we screened the remaining HC interactors using AF in order to find alternative interfaces (Figure 3A). We applied an ipTM threshold of > 0.7 to define HC direct interactors (Figure 2D). As shown, the resulting bimodal distribution may reflect AF ability to distinguish between direct and indirect binding partners.

**Figure 3:**
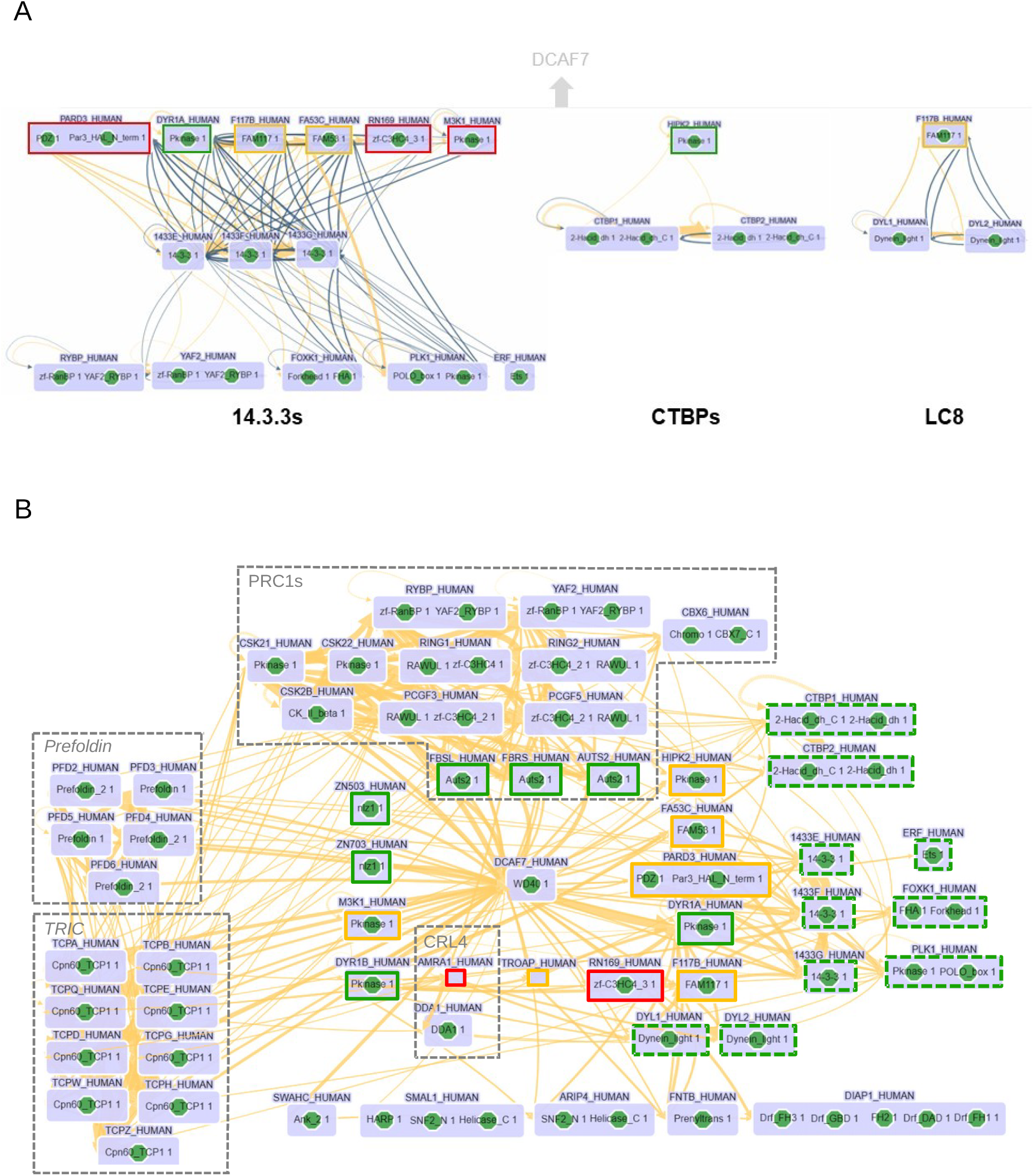
Construction of structure-based HC meta-interactome of DCAF7. **A)** Cytoscape views (SLiMAn) of SLiM-based interactors for 14-3-3 (left), CTBPs (midle) and LC8 (right). Biogrid PPIs are shown in dark yellow with edge thickness proportional to reporting frequency, IntAct PPIs in black, and SLiM-ELM PPIs are omitted for clarity. **B)** Cytoscape view of the DCAF7 HC meta-interactome. Direct interactors are outlined with solid lines colored according to the nature of the interfaces, and indirect interactors mediated via SLiMs are shown with green dashed lines. BioGRID PPIs are shown in dark yellow. Known protein complexes are outlined with dashed lines.

Fourteen proteins were predicted to bind directly to DCAF7 including the 8 previously described (DYRK1a, DYRK1b, HIPK2, ZNF703, ZNF503, AUTS2, FBRS, FBSL)(Figure 2D, colored in green). Among the newly predicted complexes, three, involving FA53C, F117B or TROAP (in orange in Figure 2D), display also linear interfaces. Their ipTMs (0.84, 0.85 and 0.74, respectively) suggested that they correspond to true direct binders through distinct SLiMs that were not further studied.

The remaining four proteins (PARD3, AMRA1, M3K1 and RNF169) are predicted to interact through domain–domain interfaces (in red in Figure 2D) according to AF predictions and their complex with DCAF7 displayed good confidence ipTM scores (0.85, 0.82, 0.75, 0.76).

Among the other proteins, 14 proteins, 5 of the prefoldin complex and 9 of the TRiC/CCT complex were considered as possible direct binders of the immature DCAF7 that might not be modeled successfully by AF (Figure 3A). Indeed, chaperonin systems such as TRiC/CCT and prefoldin mediate only transient interactions with their client proteins (Jumper et al., 2021; Baek et al., 2021; Gestaut et al., 2019). Indeed, as shown by recent cryo-EM studies, all 8 subunits of the TRiC complex cooperate for the stepwise folding of a WD40 protein (Shen and Willardson, 2025). Similarly, all the 5 subunits of the prefoldin complex appear to interact with unfolded substrates (Herranz-Montoya et al, 2021).

This analysis indicates that twenty-nine proteins out of 55 from the HC meta-interactome are likely to bind directly to DCAF7.

### Indirect binders of HC meta-interacome

Using SLiMAn, we analyzed further the DCAF7 HC interactome to identify other linear motifs from the ELM database in order to propose alternative connections to DCAF7.

For example, three 14.3.3 proteins are found (epsilon, gamma and eta) within the DCAF7 interactome, and several experimentally validated and/or high confidence phospho-motifs are present in proteins belonging to its HC meta-interactome (Figure 3A).

Experimentally validated 14-3-3 binding sites are present in DYRK1a (526-RAR<u>S</u>DP-528) and PARD3 (142-RS<u>S</u>DP-146) and several highly favorable sites are also found in 4 other direct DCAF7 partners: F117B (206-RQP<u>S</u>P-210, 270-R<u>S</u>A<u>S</u>WG-275, 343-RN<u>S</u>VE-346), FA53C (119-RSL<u>S</u>VP-124, 229-RRFSL<u>S</u>-234, 270-R<u>S</u>RSQP-275), M3K1 (17-RA<u>TS</u>P-21, 63-RKVR<u>S</u>VE-69, 260-RTVK<u>S</u>E<u>S</u>PGV-269, 1023-RKF<u>S</u>LQ-1025) and RNF169 (336-RCV<u>S</u>AP-341, 641-RQ<u>T</u>GEVGL-648) (Figure 3B). These proteins can connect indirectly the 14-3-3s to DCAF7 (Figure 3A).

As the 14-3-3s proteins are homo/hetero-dimeric they allow formation of dynamic macromolecular complexes (Yaffe et al. 1997; Howe et Barbar 2025) which can bring into a given interactome, plenty of indirect partners. Considering these properties, five other proteins: FOXK1(400-<u>R</u>GV<u>S</u>CFR<u>T</u>PF-409, 464-RYSQ<u>S</u>AP-470), ERF (466-RW<u>S</u>EDCRL-473), PLK1 (337-R<u>K</u>PL<u>T</u>VL-343), RYBP (212-R<u>SS</u>P-215) and YAF2 (166-RS<u>S</u>SP-168) with high-confidence 14-3-3 binding sites, can be indirectly connected to DCAF7. Therefore, the 14.3.3 network (Figure 3B) links indirectly to DCAF7 8 novel proteins (14.3.3E, 14.3.3F, 14.3.3G, FOXK1, ERF, PLK1, RYBP and YAF2).

In addition, two confident PXDLS-like motifs (1037-PLNLS-1041, 1145-PVSMG-1149) are present in HIPK2, a protein predicted to directly interact with DCAF7 (see above). These motifs can bring the NAD-dependent transcriptional co-repressors, CTBP1 and CTBP2, into the DCAF7 interactome (Nardini et al. 2003)(Figure 3A).

An other direct binder, F117B, contains one LC8-binding motif (246-RDKATQT-252) that can link DYL1 and 2 to DCAF7 (Howe et Barbar 2025; Jespersen et Barbar 2020)(Figure 3A).

In conclusion, SLiM-based PPIs bring 12 more proteins to DCAF7, leading to a total of 41 proteins for which molecular and structural binding modes can be delineated.

Then, we turned to dig into large assemblies to connect some of the remaining fourteen DCAF7-interactome partners still lacking a validated link.

The core subunits of mammalian PRC1 complexes are present in the DCAF7 interactome (RING1 and 2, RYBP and YAF2) (Blackledge and Klose, 2021). Other subunits including CK2 (CSK21, CSK22 and CSK2B), PCGF3, PCGF5 and the three AUTS2 homologs (AUTS2, FBRS, FBSL) characterize the mammalian PRC1.3 and PRC1.5 (Gao et al. 2012; Geng et Gao 2020), and are also present. These two particular PRC1 complexes appear to be primarily linked to DCAF7 through AUTS2, FBSL1, and FBRS, and, therefore, these three proteins may indirectly connect the proteins CK2, PCGF3, PCGF5, RING1/2,YAF2 and RYBP with DCAF7 (Figure 3C). As mentioned above, 14-3-3 proteins may play a role in these connections.

DCAF7 is also well known for its association with the CRL4 E3 ubiquitin ligase complex. Among the components of CRL the DCAF receptor AMBRA1 (also known as DCAF3) are present in the HC DCAF7 interactome (Figure 3C). However, no canonical H-box DDB1-binding motif was identified in DCAF7 (Cereghetti et al. 2008) while, AMBRA1 contains a H-box motif that mediates DDB1 binding. Consequently, DDB1 and the CRL4 would be indirectly associated with DCAF7 via AMBRA1 (Figure 3C).

Five remaining DCAF7 partners (DIAP1, ARIP4, SWAHC, SMAL1 and FTNB) do not appear to be part of well-defined stable protein complexes. Taken together, we have assigned likely binding modalities to DCAF7 for the vast majority of the proteins identified in its so-called high-confidence interactome (50 proteins out of 55) suggesting that these proteins do not correspond to spurious but rather biologically relevant hit.

### Identification of other DCAF7 direct binders

We took advantage of having constructed different signatures (regex #1 to 5) to search for potential new interactors (see Table in figure S1) within the whole DCAF7 meta-interactome (372 proteins) and the human proteome (19,413 proteins). The most stringent signatures, #1 and #2, retrieved no additional proteins. Regex 3 (reg#3, E-value = 3.7×10^−6^) retrieved no additional proteins within the DCAF7 meta-interactome, while it identified 183 proteins in the human proteome. Less stringent signatures (reg#4; E-values of 1.0×10^−4^ and reg#5; 1.5×10^−5^) retrieve 42 and 14 proteins in the DCAF7 meta-interactome, respectively, and 1240 and 402 in the human proteome. For each hit (beside the hits of reg#4 in the human proteome), we modeled the putative complexes with DCAF7 using AF. Then, we evaluated their ipTM scores corresponding to models for the FL proteins as well as for the 50-residue motif-centered fragment (Frag50)(Figure S1). We considered as potential valuable new interactors the proteins for which the ipTM of the heterodimer AF model is better than 0.7 for either Frag50 or FL (Figure S1). We did not consider further proteins for which the interface of the frag50 is not centered. No hits were found with similar properties (similar interface as FL and Frag50 with high AF scoring) as the most confident binding partners (circles). Twelve proteins (in yellow in Figure S1) show high ipTM in the FL protein configuration, but low ipTM for their Frag50, while showing a linear interface with DCAF7. In parallel, 17 proteins show domain–domain interfaces with DCAF7 with high AF ipTM score predictions in the FL configuration (in red in Figure S1). Finally, 6 proteins displayed accurate AF models for the Frag50 segments despite unfavorable modeling in their FL configuration (pale green).

Representative examples from the three different categories of the newly predicted heterodimers are presented in figure 4. Binding of DDX41 and MED13L is predicted to occur through the DCAF7 SLiM motif that docks onto the top surface of the DCAF7 β-propeller (Figure 4A). Other interactors bind to DCAF7 also via distinct linear motifs and on the side of the β-propeller (LZTS3, SHISA6)(Figure 4B). Finally, a third category like for EHMT2 involves domain–domain interfaces (colored in black) (Figure 4C). Overall, through these extensive searches and subsequent modeling studies, we identified 35 potential new direct interactors, including 9 within the whole DCAF7 meta-interactome and 26 within the human proteome.

**Figure 4:**
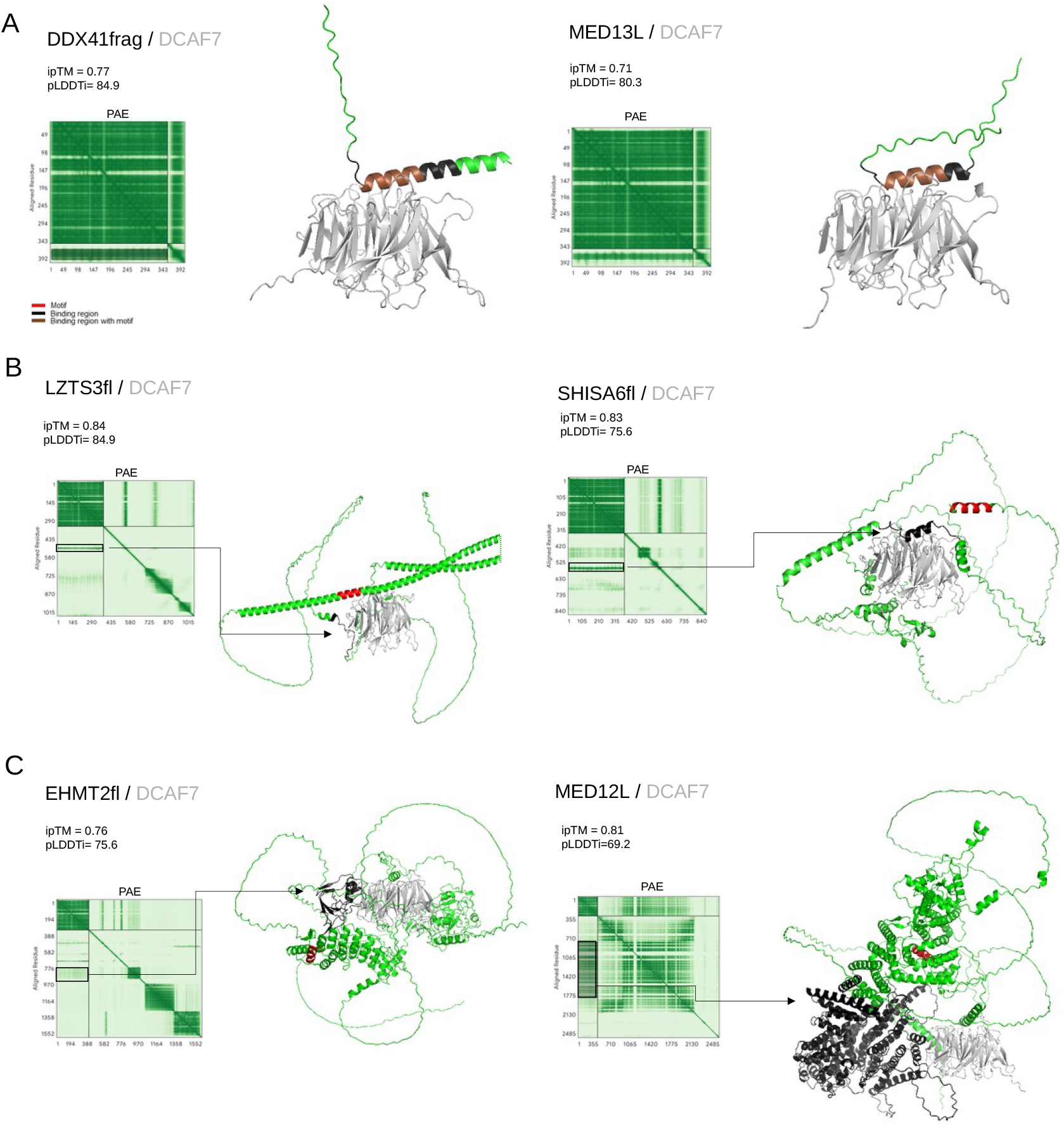
Identification of potential direct interactors of DCAF7 in its meta-interactome and the human proteome. Examples of AF models illustrating the different types of binding interfaces identified in the retrieved hits (sup.figure 1): **A)** DCAF7 motif ; **B)** other linear interfaces ; **C)** domain-domain. For each interface, the ipTM score, pLDDT of the interface residues (pLDDTi), and the PAE plot are shown. The binding regions delineated in the PAE plot (arrow) are colored as indicated.

### Digging into the interconnectivity of interactors

In an attempt to validate all our hits, we defined an interconnectivity score for the DCAF7 potential partners based on two criteria: the number of shared interactors between the DCAF7 meta-interactome and that of the the protein of interest, and the robustness of the corresponding PPI (i.e., the number of times the PPI was observed). The interconnectivty score (a Jaccard score described in the Material and Methods section) was defined by a simple heuristic where zero means no common partners and the maximum theoretical score is one, corresponding to identical meta-interactomes between the two proteins. Higher scores reflect a greater proportion of shared interactors. For the 8 partners of the HC-Meta-interactome with the newly defined SLiM-based interaction, the score ranges from 0.16 to 0.82 with a mean score of 0.51 (Figure 5). High scores were obtained for the three paralogs, FBRS (0.82), AUTS2 (0.70) and FBSL (0.61) and putatively reflecting their stable association with DCAF7 within the major transcriptional regulator PRC1 complex (Figure 5). DYRK1A and B kinases, that are also very frequently associated with DCAF7, also show strong interconnectivity scores of 0.77 and 0.62 respectively (figure 5). For the potential direct interactors in the DCAF7 LC-Meta-interactome, the mean interconnectivity score is lower (0.3), with M3K1 showing the highest score (0.5).

**Figure 5:**
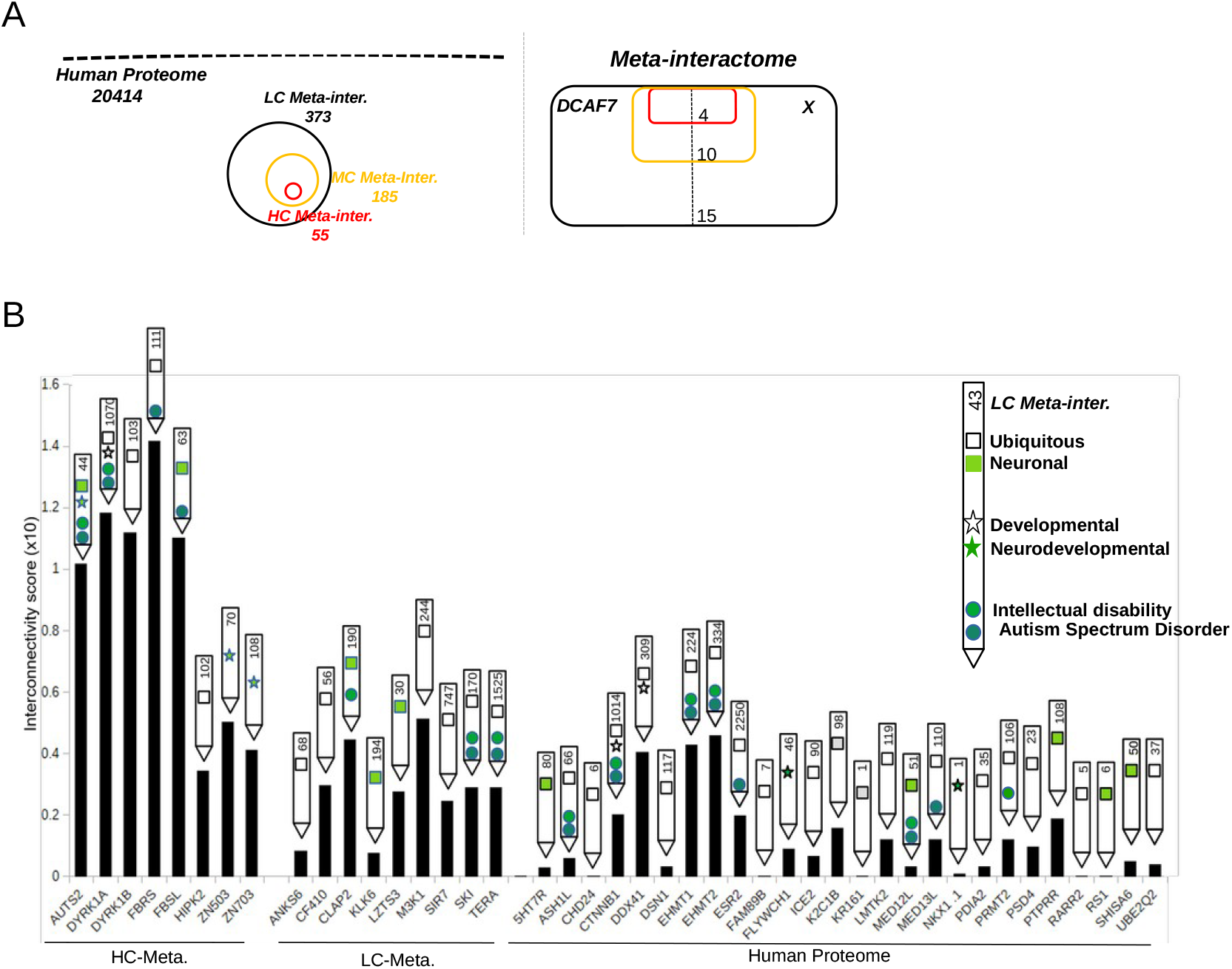
Interconnectivity of DCAF7 interactors. **A)** Schematic representation illustrate the size (compare to human proteome) of meta-interactomes (DCAF7 as example) with different confident level (H: high, M:medium, L:low confeidence) (left). On the right, selection of common interactors in each meta-interactome used for interconnectivity scoring. B) Interconnectivity score for each direct binders. Legend indicates the number of proteins in each meta-interactome, as well as the tissue and temporal expression, and the associated diseases for each partner.

In the case of the 26 potential new DCAF7 interactors from the human proteome, 5 proteins have a null interconnectivity with DCAF7 whereas 21 proteins have a positive interconnectivity score with a mean of 0.14 (ranging from 0.04 to 0.45) that require additional evaluation to be consider as probable direct binders. However, three proteins (DDX41, EHMT1, and EHMT2) have interconnectivity scores (0.40, 0.43, and 0.46) comparable to those of some direct binders (i.e., HIPK2, ZNF503, and ZNF703), making them good candidates for direct PPIs with DCAF7 (Figure 5A). (Figure 5A). The other putative partners showed rather weak interconnectivity and for most of them it might not be sufficient per se to validate their belonging to the DCAF interactome. Nevertheless, among them, we can observe the presence of two subunits of the mediator complex, MED12L (0.03) and MED13L (0.12), that are associated to the MED23 subunit which is present in the DCAF7 LC meta-interactome. Of note, two additional mediator subunits, MED12 and MED13, are identified in the human proteome (with the Regex 3 and 5, respectively). MED13 is at the limit of our AF selection threshold, with ipTM scores of 0.7 for the FL model and 0.6 for Frag50, whereas MED12 shows modest ipTM scores of 0.43 for the FL model and 0.34 for the Frag50 model. Another protein, the beta-catenin (CTNNB1) that has been known to interact with DYRK1A appears to be another possible binder of DCAF7 although the experimental evidence (Liu et al. 2022) is not listed in BioGrid or Intact databases. Taken together, these results support the use of such interconnectivities to extend further the meta-interactome of DCAF7.

### Biological relevant correlation among DCAF7 interactors

We have also noted a strong overlap between the tissular and temporal expression of DCAF7 and its partners, with principally two categories corresponding to ubiquitous or neuronal expressions. Furthermore, one third of these proteins are expressed during neuronal development (Figure 5). We observe also a strong enrichment for genes involved in neurodevelopmental diseases such as autism spectrum disorder (ASD), intellectual disability (ID) (Figure 5). In fact, the current experimental DCAF7 meta-interactome (374) contains 44 genes linked to autism and annotated in the SFARI Gene database revealing a significant enrichment (1.97, p = 1.3 × 10^−5^). Furthermore, higher enrichment is observed for the direct interactors containing a DCAF7 motif (6/17) within the DCAF7 meta-interactome (5.88, p = 3.2 × 10^−4^) (Figure 5). Interestingly, high significant enrichment (4.76, p = 1.5 × 10^−3^) for autism-related genes is also found for the potential new direct binders (7/21). The two mediator subunits MED12L and MED13L are linked to autism like MED23 and MED13. Also, highly relevant for ASD, are the neuronal protein SHISA6, which interacts with and modulates AMPA receptors, as well as DDX41, which interacts with and regulates phosphorylation of SRRM2, a protein linked to autism as well (Klaassen et al. 2016; Regan-Fendt et al. 2023)

In summary, these potential new interactors were selected based on the presence of the motif together with high-confidence AF models and the observed enrichment in autism-associated genes further supports their relevance as *bona fide* DCAF7 binders. This strong link to ASD correlates with the high interconnectivity of DCAF7 with high-risk autism genes, DYRK1A, AUTS2, and FBSL1 (Fu et al. 2022).

### Extended putative interactomes

To strengthen the relevance of the above predictions, we modeled several oligomeric complexes possibly linked to neuronal signaling and autism (Figure 6). For example, known SLiM-based PPIs show a link between LZTS3 and SHANK3 via its PDZ domain of, as well as the presence of several DYRK1A/B phosphosites in SHANK3 (T1204, S1309, S1711, S1713, S1715). AF modeling of the heterotetrameric assembly showed high scores for binding interfaces between DCAF7–DYRK1A and DCAF7–LZTS3 that are similar to that of in the corresponding heterodimeric complexes described above (Figure 6A). We were also able to model a heterotetrameric complex composed of DCAF7, DYRK1A, LZTS3, and SHANK3 PDZ domain with high accuracy (Figure 6A). The kinked α-helix between DCAF7-DYRK1A in the heterotetramer is positioned similarly that in the heterodimer and involved the DCAF7 binding motif in DYRK1A. The C-terminal PDZ-binding site (668-IESTEI-673) of LZTS3 is also correctly positioned within the groove of the PDZ domain of SHANK3 (Figure 6A). Interestingly, two ASD-linked mutations in SHANK3 (R1708H, P1714L) destroy DYRK1s phosphosites while other missensse variants have been also found in the PDZ domain (Waga et al. 2011). Therefore, our model supports a functionally important phosphorylation of SHANK3 by DYRK1A that would be assisted by two scaffolding proteins, DCAF7 and LZTS3.

**Figure 6:**
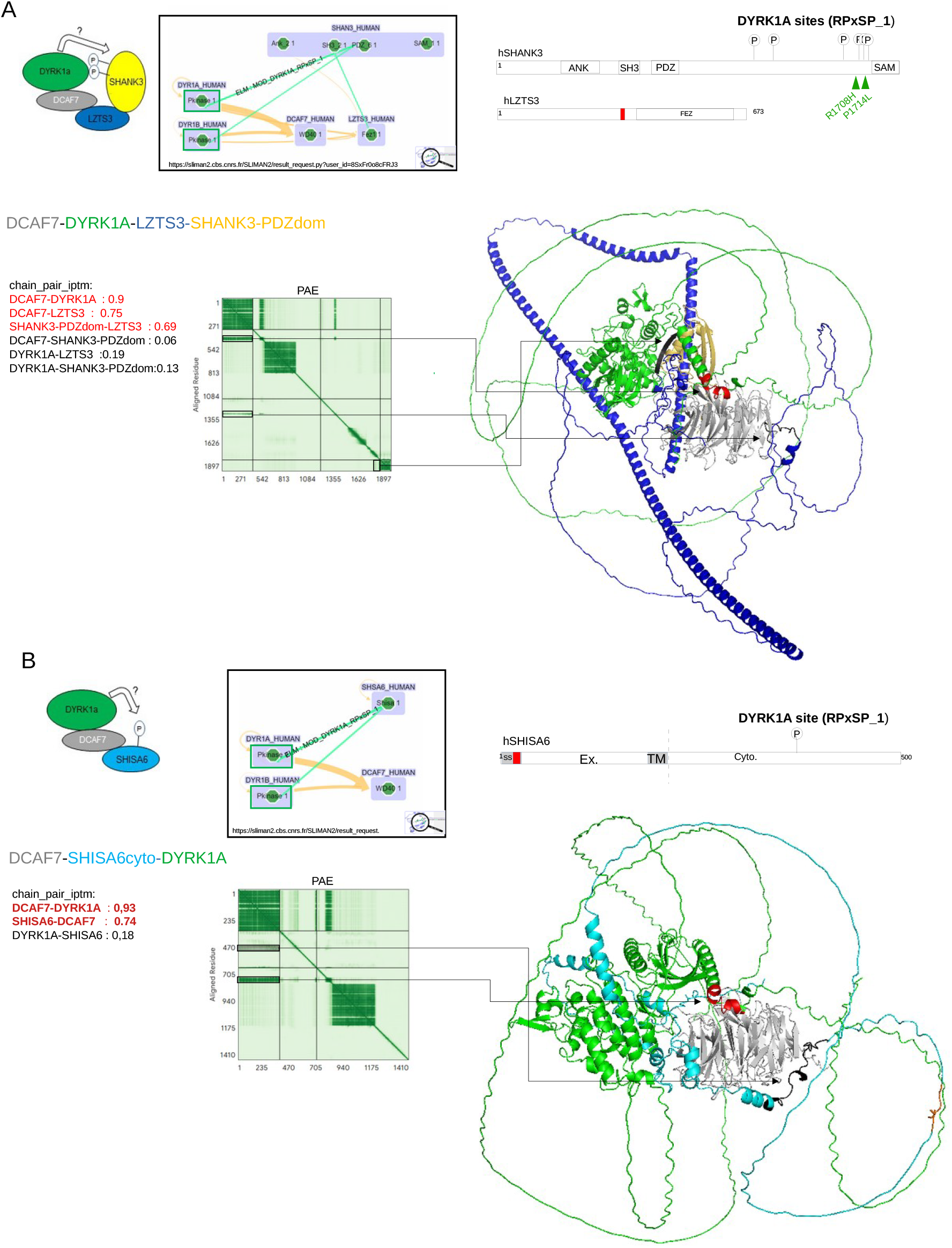
AF models of functionally relevant protein complexes. **A)** Role of LTZ3 in the DYRK1A/B kinase module mediating SHANK3 phosphorylation and **B)** Phosphorylation of SHISA6 by DYRK1A/B kinases module The cytoscape views of the oligomers were generated by SLiMAn. BioGRID PPIs (dark yellow) and DYRK1A/B phospho-SLiM entries (green lines) are indicated. AF models shown in PyMOL cartoon view, with proteins colored as indicated. Pairwise interface confidence scores (ipTM) are shown for each interacting chain pair in the oligomeric complex. Functional model of oligomeric complexes are also presented.

Another complex linked to neuronal signaling and ASD can be built with SHISA6, an auxilary subunits of AMPA receptors at the postsynapic membrane (Figure 6B). As highlighted by SLiMAn, a relevant DYRK1 phosphorylation site is present in the cytoplasmic region of SHISA6 (serine S373) (Figure 6B). The AF model of the heterotrimer DCAF7–DYRK1A–SHISA6 reproduces the interfaces previously predicted in the DCAF7-DYRK1A and DCAF7–SHISA6 heterodimers. No relevant interface (chain_pair_ipTM = 0.18) was observed between DYRK1A and SHISA6 (Figure 6B). Like for SHANK3, this model supports a possible phosphorylation of SHISA6 by DYRK1A or/and DYRK1B. Taken together, these models illustrate how DCAF7 can bring the DYRK1 kinases to the postsynaptic proteins SHANK3 and SHISA6, and drives their phosphorylation. This highlights a mechanistic link between DYRK1-mediated signaling at excitatory postsynaptic compartment, that seems highly relevant to autism.

## Discussion

Our integrative structural and interactomic analysis provides compelling insights into the molecular architecture of DCAF7. Using AF structural modeling, in combination with motif analysis via SLiMAn, we identified and validated key PPIs interfaces, particularly on the conserved top face of the DCAF7 β-propeller. This surface emerges as the central pocket for direct binding events, highlighting its functional importance across distinct cellular pathways. The structural modeling revealed that DCAF7 recognizes a defined subset of protein partners through a SLiM encompassing an important α-helix observed in disordered region of DYRK1AB kinases. Interestingly, this specific interface, supported by co-immunoprecipitation and mutagenesis data, appears to be conserved across another kinase, HIPK2, suggesting a shared binding mechanism within a functionally diverse set of DCAF7 partners. This will support the rule of DCAF7 in the coordination of multiple kinase activities.

Similar molecular features appear to be involved in the direct binding of DCAF7 to the transcription repressors ZNF503 and ZNF703. Interestingly, DCAF7 is known to be present with ZNF703 in a nuclear transcriptional repressor complex composed of the prohibitin (PHB2) protein and NCOR2 (Sircoulomb et al. 2011). The role of DCAF7 is unknown but this repressing complexe is involve in the transcriptional hypoactivity of eostrogen receptors in breast cancer cells (Sircoulomb et al. 2011). Of note, both ZNF703 and DCAF7 appears upregulated in luminal B breast cancers (Sircoulomb et al. 2011). Also involved in breast cancer aggressiveness, but through a distinct mechanism, ZNF503 has been found to be associated with DCAF7, principally in high-throughput screening.

The very same DCAF7 pocket at the top of the b propeller appears to be also involved in the binding of the three paralogs AUTS2, FBSL1 and FBRS (figure 2). DCAF7 is often found associated with these proteins and the binding region have been map for AUTS2 (600-700), that contain the DCAF7 motif we defined above (Wang et al. 2018). These paralogs are present in PRC1.3 and PRC1.5 complexes (Hauri et al. 2016; Gao et al. 2012). In accordance with the direct interaction with these subunits, DCAF7 is also found uniquely in PRC1.3 and PRC1.5 (Gao et al. 2012). While poorly investigated, so far, the role of DCAF7 in PRC1 complexes appear essential for transcriptional activity of non-canonical PRC1s (Wang et al. 2018).

In addition, we have partially elucidated the scaffolding mechanism of DCAF7, involved in the coordination of the three kinases DYRK1A, HIPK2, and M3K1, described previously by Ritterhoff et al. (Ritterhoff et al. 2010). The fact that HIPK2 has two modalities of binding to DCAF7, via a SLiM and by domain-domain interfaces, suggests that DCAF7 can scaffold DYRK1A and HIPK2. Similar scaffolding can be envisaged in the case of M3K1, which appears to bind preferentially DCAF7 via a domain-domain interface but, like HIPK2 also contains a DCAF7 binding motif. In the same line, DCAF7 connects the adenoviral protein E1A with DYRK1A and HIPK2, leading to its subsequent phosphorylation (Glenewinkel et al. 2016; Ritterhoff et al. 2010). We have structure modelling evidences on the scaffolding properties of DCAF7 with DYRK1A, HIPK2, and E1A. The binding interface of E1A to DCAF7 is close to what we observed for several other partners (SKI, LZTS3, LZTS2, SHISA6), with linear motifs interacting on the side of the β-propeller (Figures 4 and 5).

In this study, we highlighted the identification of several new PPIs interfaces for highly relevant ASD-linked proteins. Although DCAF7 is not *per se* an autism risk gene, likely due to strong evolutionary pressure acting on this essential gene, 12 proteins (DYRK1A, AUTS2, SKI, SYNGAP1, CUL3, NIPBL, MED23, MED13L, ASH1L, EHMT1, CTNNB1, and SHANK3) present in its extended high-confidence meta-interactome, do belong to the 72 most confident high-risk autism genes (Fu et al. 2022). Furthermore, we have found strong structural and interconnectivity evidences in favor of direct interactions between high-risk autism-linked proteins (DYRK1A, AUTS2, SKI, MED13L, ASHL1, EHMT1 as well as CTNNB1) and DCAF7.

ASD is caracterized by a high heterogeneity in genetics and non genetics factors, that ultimately converge to synaptic defects (Engal et al. 2024). Approximately one-third of the 1200 autism-linked genes encodes proteins that directly participitate in synaptic function at both presynaptic and postsynaptic compartments (Sauer et al. 2026). Among the proteins most frequently associated with DCAF7, the kinase DYRK1A displays a widespread neuronal localization, encompassing synaptic compartments as well as the nucleus (Atas-Ozcan et al. 2021). As is typical for protein kinases with numerous substrates, loss of DYRK1A function in neurons results in widespread effects on multiple targets, including Tau, NMDA receptors, ABLIM3 and β-tubulin (Atas-Ozcan et al. 2021). Here, we present high confident structural models showing that DYRK1A may indirectly interact via LZTS3, and and phosphorylate the postsynaptic protein SHANK3 (Figure 5). Similarly (AF model with high confident score: 0.82) can also be obtained with another autism-linked protein, LZTS2, a close homolog of LZTS3. Interestingly, ASD-linked rare variants in SHANK3 appear to affect DYRK1A phosphosites located at its C-terminus (Figure 5). We also identified another probable postsynaptic substrate of DYRK1A, the auxiliary subunit of AMPA receptors, SHISA6. The latter modulates AMPA receptors activity and synaptic transmission (Klaassen et al. 2016; Peter et al. 2020). These two DYRK1A substrates directly modulate key components of excitatory glutamatergic synaptic transmission providing mechanistic insight into genotype–phenotype relationships in ASD (Bonsi et al. 2022).

Several genes linked to ASD are implicated in epigenetic and transcriptional regulation (Engal et al. 2024). AUTS2, an other direct interactor of DCAF7, is also linked to alterations in ASD (Pang et al. 2021). Interinstingly, we isolated in the human proteome, three subunits of the mediator complex (MED12L MED13L and MED13) that are also linked to ASD. The mediator complex composed of twenty-six subunit is a master regulator of RNA polymerase 2 (Richter et al. 2022). The above three subunits are part of the Mediator kinase module and bind also CTNNB1 (Kim et al. 2006). Taken together, we found that 53 proteins linked to ASD are direct or indirect partners of DCAF7/ All these results point to an important but so far hidden role of DCAF7 in ASD.

Our study has highlighted different protein–protein interaction (PPI) interfaces of DCAF7 with its numerous partners and identified several potential additional partners for which no studies have yet been reported. Based on various structural models and interactomics data, we have deciphered the principles of DCAF7 scaffolding mechanisms that link some important protein-kinases to their substrates. In conclusion, DCAF7 emerges as a regulator of kinase activities, enabling interactions between various cellular components, particularly those involved in postsynaptic signaling linking them to neurodevelopmental disorders.

## MATERIAL AND METHODS

### Interactome analysis

Protein interactors were retrieved from BioGRID (https://thebiogrid.org) and IntAct (https://www.Qebi.ac.uk/intact/) and analyzed using SLiMAn (http://sliman.cbs.cnrs.fr). SLiM motif selection using SLiMAn 2.0 (https://sliman2.cbs.cnrs.fr) was performed as previously described (Mezghrani et al. 2024). Regular expressions (regex) were generated using PSSMSearch (https://slim-tools.org/tools/pssmsearch/input) and subsequently used to identify potential new interactors with SLiMSearch4 (https://slim-tools.org/tools/slimsearch/input).

### AF structural modeling

Protein complexes were modeled using AF (in-house AlphaFold2 multimer and AlphaFold3 (https://alphafoldserver.com). Protein sequences used for AF modeling were retrieved from UniProt (https://www.uniprot.org). For motif-based interactions, 30- or 50-residue fragments centered on the motifs were extracted. AF models with an ipTM score ≥ 0.8 were selected for motif extraction, while models with ipTM ≥ 0.7 were considered as valid complexes. Three independent AF simulations were performed for each retained model, and the most reliable model was selected for downstream analyses. Residue-level confidence was assessed using pLDDT scores, and predicted aligned error (PAE) values were also taken into account to evaluate the reliability of the predicted interfaces. Motifs were defined by superimposing the binding interfaces, and residues occupying equivalent positions across models were retained for motif definition.

### Interface analysis of protein multimer and structural visualization

Structural models were analyzed and visualized using PyMOL (https://www.pymol.org). Structural superimpositions and RMSD calculations were also performed in PyMOL.

### Interconnectivity score

A dedicated interconnectivity score was defined to quantify the connectivity between proteins. For each protein pair (e.g., DCAF7 and protein X), the number of interactors common to both proteins was determined. These common interactors were categorized with SLIiMAn as observed at least twice in BioGRID and IntAct (bio2, int2), observed only once in either database (bio1,int1), or included in the total. The counts in each category were then used to calculate the interconnectivity score as follow: IntS(A,B)=((Meta(A)bio2int2 ∩Meta(B)bio2int2)/(Meta(A)bio2int2∪Meta(B)bio2int2))+((Meta(A)bio1int1∩Meta(B)bio1int1)/(Meta(A)bio1int1∪Meta(B)bio1int1))+(Meta(A)∩Meta(B))/ (Meta(A)∪Meta(B)))/3.

## Legends

**Supplementary Table 1.**
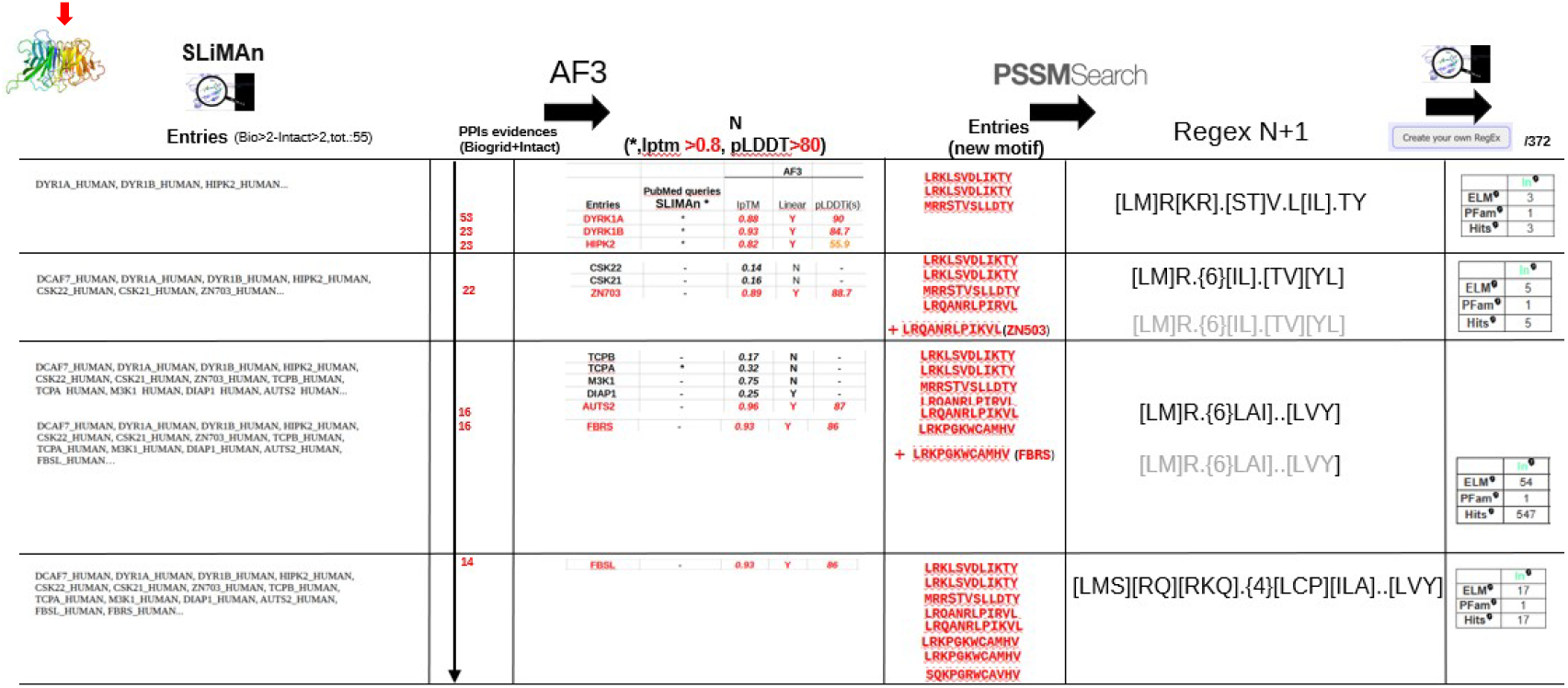

**Supplementary Figure 1.**
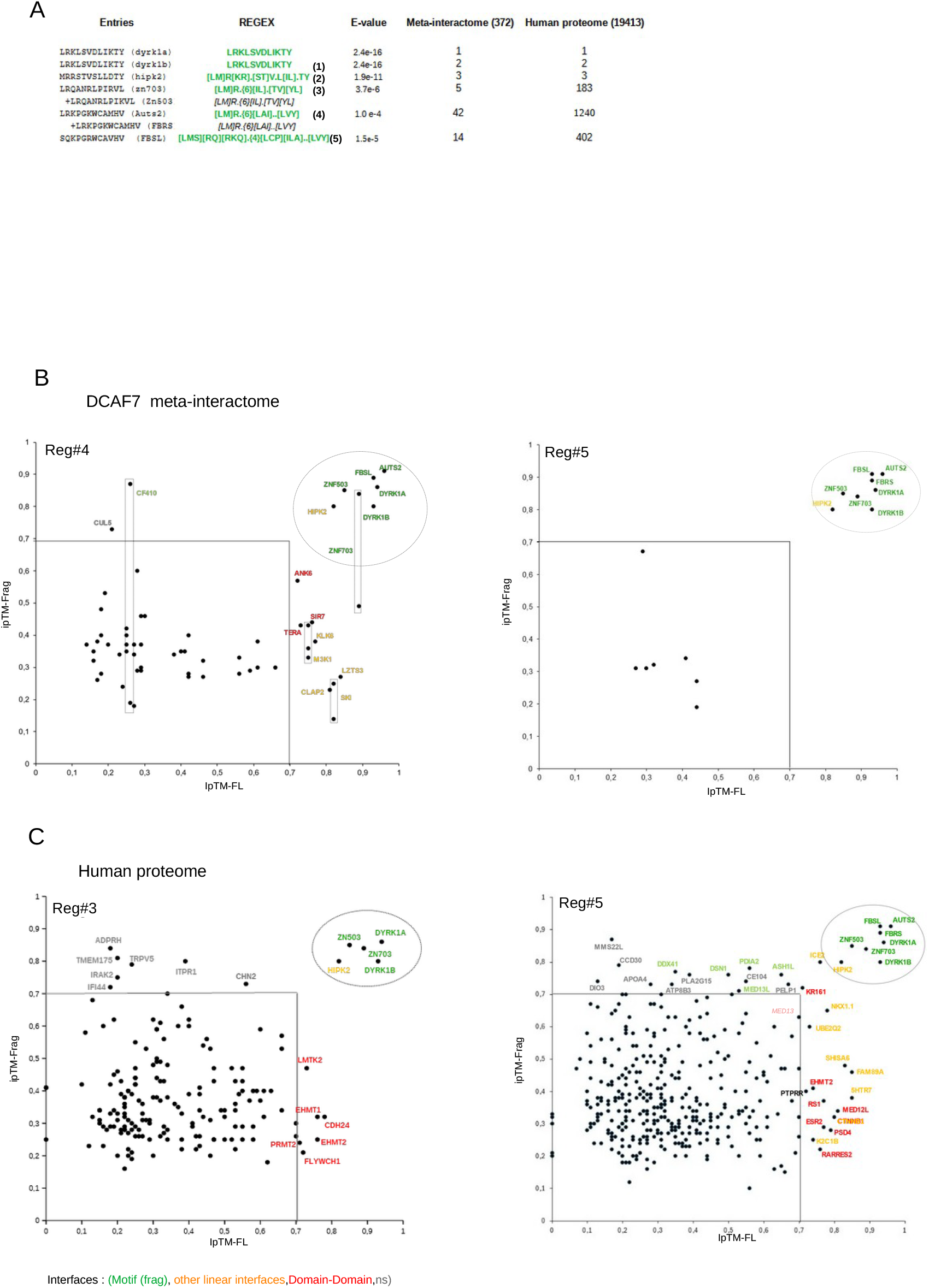
**A)** Table presenting the generation of different regex with PSSMSearch, their E-values, and the number of hits in the DCAF7 meta-interactome and the human proteome. **B-C)** 2D plots of ipTM scores obtained for AF models of 50-residue fragments containing motifs (ipTM-Frag) as a function of FL proteins (ipTM-FL) for each hit. Hits were obtained using regex (Reg.) as indicated in the DCAF7 meta-interactome (B) and in the human proteome (C).

